# ContrastQA: A label-guided graph contrastive learning-based approach for protein complex structure quality assessment

**DOI:** 10.1101/2025.06.20.660832

**Authors:** Lei Zhang, Rui Ding, Xiao Chen, Jie Hou, Dong Si, Yang Wang, Keying Lin, Renzhi Cao

## Abstract

Despite recent progress, the Estimation of Model Accuracy (EMA) for protein complexes remains less advanced compared to that for protein monomers. A key challenge lies in effectively integrating both interface-specific and global structural information to accurately assess the quality of protein complexes. Here, we introduce ContrastQA, the first EMA framework for protein complexes that incorporates the proposed label-guided graph contrastive learning based on interface quality. By integrating a geometric graph neural network to model global structural features, ContrastQA effectively captures both local (interface-level) and global (structure-level) information for accurate model quality estimation. ContrastQA achieved ranking losses of 0.123 and 0.116 on the TMscore and GDT-TS metrics on the CASP16 dataset, which are 0.015 (10.9%) and 0.012 (8.7%) lower than the second-best EMA method with ranking losses of 0.138 and 0.128. Furthermore, for the more challenging CASP16 hard targets, our method achieved a TMscore ranking loss of 0.131, which is 0.023 (14.9%) lower than that of the second-best EMA method with a ranking loss of 0.154. Our study demonstrates the strong effectiveness of the label-guided graph contrastive learning module. These findings suggest that our graph contrastive learning framework serves as a valuable pre-training strategy for learning protein structure representations. The source code is available at https://github.com/Cao-Labs/ContrastQA.

**Significance Statement:** High-quality proteins generated by high-precision structural prediction methods for protein complexes helping biologists prioritize models for drug discovery and understanding disease mechanisms. Recent advances in predicting protein complex structures (like AlphaFold3) still struggle to match experimental precision. A critical bottleneck lies in accurately estimating the quality of predicted models-known as Estimation of Model Accuracy (EMA). Here, we introduce ContrastQA, implementing label-guided graph contrastive learning framework to explore the generalization of high-quality proteins. Our method demonstrates outstanding performance, provides an effective solution for assessing the structural quality of protein complexes.

---

**P**rotein-protein complexes have emerged as central components in elucidating biological functions and cellular processes. With the advent of deep learning-based prediction methods such as AlphaFold3 (1) and RoseTTAFold All-Atom (2), the precision of protein complex structure prediction has approached that of traditional conformational sampling approaches. However, unlike protein monomer prediction (3), the accuracy of protein multimer prediction remains suboptimal, so improving the accuracy of protein complex prediction is an important and challenging issue. Consequently, Estimation of Model Accuracy (EMA) for protein complex structures has garnered significant attention as a critical component of protein complex structure prediction pipelines. Therefore, EMA for protein complex structure was introduced as a new evaluation category in CASP15 (4–6), reflecting its growing importance in computational structural biology.

Specifically, EMA methods aim to score from a pool of decoys for the purpose of improving the accuracy of protein complex structure’s prediction. Current EMA approaches are broadly categorized into two groups: multi-model methods and single-model methods. As the name implies, multi-model approaches, such as MULTICOM_GATE (7), ModFOLDdock2 (8), and VoroIF-jury (9), leverage structural alignments and consensus strategies to evaluate model quality. While effective, their performance is inherently constrained by the diversity and quality of the input model pool. In contrast, single-model methods, including GNN-DOVE (10), ComplexQA (11), EnQA (12), DProQA (13), VoroIF-GNN (14), GraphCPLMQA (15) and TopoQA (16), et.al. They require only a single predicted structure as input. All of these methods leverage protein sequences, physicochemical properties, and geometric structural information to learn the relationship between individual features and structure quality using deep learning techniques. Notably, recent advancements in single-model EMA methods have demonstrated remarkable progress, even surpassing the performance of traditional multi-model approaches in several key aspects. The single-model EMA methods had been comparable to the multi-model EMA methods in terms of interface quality evaluation of protein complexes in CASP15 (4). Therefore, single-model EMA methods are bound to become popular in the field of protein structure quality accuracy estimation. Recently, the category of EMA for complex structures and their interfaces introduced by the latest Critical Assessment of Techniques for Protein Structure Prediction (CASP) competition — **CASP16** assesses three main aspects: Topology Global Score (TGS), Interface Total Score (ITS) and Interface Residue-Wise Score (IRWS).

Protein complexes consist of multiple monomers, with protein-protein interfaces—the contact regions between monomers—playing a crucial role in mediating interactions such as signaling and metabolism (17). Given their functional importance, many existing methods focus on assessing interface quality, making the accurate representation and effective learning of interfacial features a central challenge (11, 14, 16). There are also many methods to assess the global structure quality of protein complexes, adapting features from monomer-based EMA to protein multimers and designing features specific to the properties of multimers (7, 13, 15). However, effectively integrating interface information with global structural features remains a key challenge. The global EMA methods for protein multimers mainly focused on the overall characteristics of the complexes and the features of the monomers of the internal components. If interface information can be effectively exploited, leveraging local structural cues to optimize global quality estimation may significantly enhance the precision of protein complex scoring. Current approaches typically feed features extracted from protein sequences and structures directly into deep learning models to learn embedding representations. However, this direct strategy often results in suboptimal learning performance, as much of the informative content is lost during the propagation process within the network.

To address this issue, we draw inspiration from MULTICOM_GATE (7), which observed that high-quality decoys under the same target often exhibit structural similarity, whereas low-quality ones tend to vary significantly. This is highly consistent with the main idea of contrastive learning, which has emerged as a powerful technique for learning robust and discriminative graph representations (18–20). It aims to construct a feature space in which similar (positive) samples are pulled closer and dissimilar (negative) samples are pushed apart, thereby enhancing the model’s ability to distinguish between structural variations. However, most conventional contrastive learning approaches are unsupervised and rely heavily on data augmentation or perturbation to define positive and negative pairs (19–21), without considering domain-specific structural knowledge. This observation inspired us to rethink the sample pairing strategy in the context of EMA for protein complexes. Based on MULTICOM_GATE we delved into the composition of protein complexes in an attempt to incorporate interface information into contrastive learning and extend it to a single-model approach. By incorporating domain-specific supervision — particularly the interface quality scores such as DockQ(22) — we designed a label-guided contrastive learning framework tailored to the protein complex EMA task. This approach utilizes the interface-based labels to guide the construction of contrastive pairs, enabling the model to more effectively capture both local (interface-level) and global (structure-level) features, and ultimately improving protein graph embedding learning.

Here, as shown in Fig. 1A, we present **ContrastQA**, the first single-model method for EMA of protein complexes based on a novel graph contrastive learning framework. The model combined geometric graph neural networks (GVP-GNN) (23) with contrastive learning for protein structure features learning. A key contribution of ContrastQA is a task-specific strategy for constructing positive and negative samples pairs, guided by the interface quality metric DockQ (22, 24), as shown in Fig. 1B and Fig. 1C. Specifically, samples were categorized as positive or negative based on their DockQ scores, and combined with anchor samples to form groups. This targeted pairing enables the model to learn rich, high-order local residue-level structural representations. We also designed a contrast loss function adapted to our strategy, which added labeling information to the traditional contrast loss function and utilized the labeling information to enhance the labeling guidance. Furthermore, we adopted a geometric graph neural network architecture (23) that leverages rotational invariance, allowing the integration of 1D and 2D features with 3D geometric information in a physically meaningful way. More specific details can be found in the section Materials and Methods 1.

**Fig. 1.**
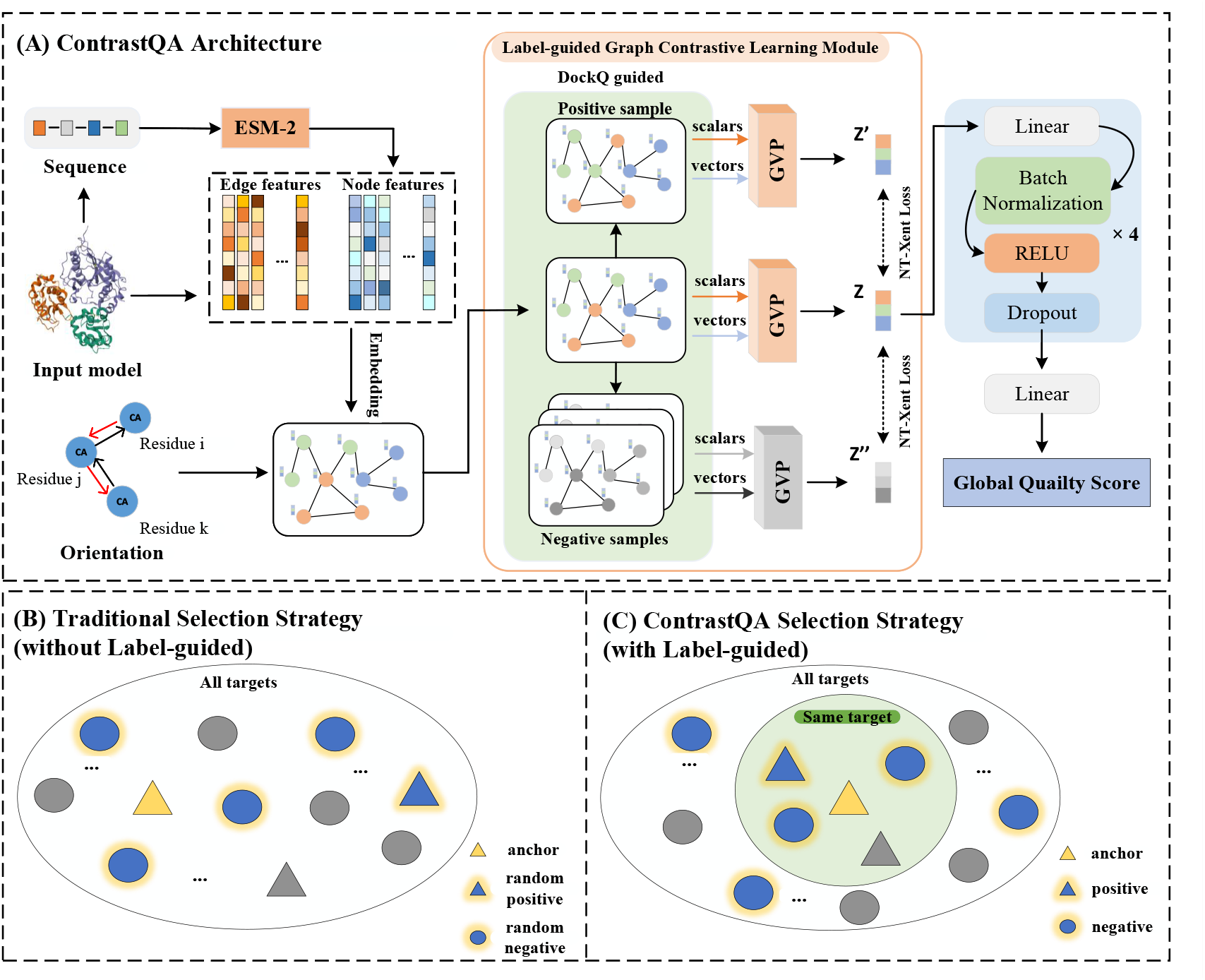
**(A)** Overview of ContrastQA. The global quality score of a protein structure is predicted using a label-guided graph contrastive learning module combined with a geometric graph neural network. Scalar and vector features extracted from the protein complex are embedded into a KNN graph, which is then input into a label-guided graph contrastive learning module, where positive-negative samples pairs are defined based on interface metric DockQ score labels. The resulting feature embeddings are passed through a multilayer perceptron (MLP) to produce the final global quality score. **(B)** and **(C)** Positive-Negative Sample Selection Strategies in Traditional Methods vs. ContrastQA. (B) Traditional methods randomly select negative samples (blue circles) from the entire sample space, without considering target identity or structural similarity. (C) ContrastQA introduces a multi-granularity sample selection strategy. For each anchor sample (yellow triangle), positive samples (blue triangles) and negative samples (blue circles) are first selected from decoys belonging to the same target. If insufficient negative samples are found, additional ones are drawn from the broader sample space. Additionally, DockQ score labels are employed to guide the selection process, enabling more precise distinction between positive and negative samples.

## Results

### Evaluation metrics

We used the constructed protein complex structure dataset to train the model and used it to test non-redundant CASP protein multimers as well as multimers composed of structures predicted by AF3. In the tests, we assessed the global quality of the multimers. For global quality assessment, we employed the *Top1 loss* (i.e., ranking loss) based on TMscore (25) and GDT-TS (26). TMscore evaluates the global similarity between the predicted and native protein structures and is highly robust to variations in protein size and shape. The TMscore ranges from 0 to 1, with values closer to 1 indicating higher structural similarity. We used US-align (27) to compute the TMscore. In contrast, GDT-TS focuses on local structural matching by measuring the proportion of residues that align within predefined distance thresholds, without applying global normalization. Like TMscore, GDT-TS also ranges from 0 to 1, with higher values indicating greater similarity. We utilized Openstructure (28) to compute GDT-TS. *Top1 loss* is defined as the difference between the true score of the predicted top-ranked decoy and that of the actual top-performing decoy. A smaller value indicates higher prediction accuracy of the method.

### Baselines

To comprehensively evaluate the performance of ContrastQA, we compared the results with some of the best deep learning-based methods in recent years: GNN-DOVE (10), ComplexQA (11), DProQA (13), VoloIF-GNN (14), and TopoQA (16). GNN-DOVE is a method for assessing the structure of protein complexes using gated-enhanced GAT, and ComplexQA is a method for assessing the structural quality of protein complexes using a combination of GCN and a wide range of physicochemical features and characterization techniques for amino acids. DProQA and VoroIF-GNN are two of the top-performing single-model methods for EMA in CASP15. TopoQA is the latest method to assess the structural quality of protein complexes, which introduces topological features from topological deep learning and effectively combines GAT.

### Performance on the CASP16 dataset

The CASP16 dataset is from the Protein Multimers section of the latest CASP competition. We tested ContrastQA on 37 targets on the CASP16 dataset and compared it to multiple deep learning-based methods for EMA of protein complexes.

As shown in Table 1, we compared the TMscore *Top1 loss*, the GDT-TS *Top1 loss* on CASP16 with a variety of advanced methods. We also compared it with the state-of-the-art (SOTA) methods in CASP16, which are multi-model as well as single-model methods. Since most of the methods do not have published code, we made the comparison directly based on the data published on the official CASP16 website. In addition, since the native structures of targets H1229 and H1230 used by the CASP teams were not publicly available, and targets H1265, T1219o, and T1295o were excluded from the original result statistics, we standardized the target selection for all comparison methods. Specifically, targets H1229 and H1230 were removed and replaced with H1265, T1219o, and T1295o to ensure fairness in result comparison. The CASP teams also evaluated the latter three, and we used the evaluation results, then we utilized US-align and Openstructure to calculate the TMscore and GDT-TS scores, respectively, to obtain the final ranking loss results. It can be seen that our method achieved the best results in terms of *Top1 loss*.

**Table 1.** Performance on the CASP16 dataset.

| Method | Top1 loss |  |
| --- | --- | --- |
|  | TMscore (↓) | GDT-TS (↓) |
| GNN-DOVE | 0.332 | 0.315 |
| DProQA | 0.213 | 0.275 |
| ComplexQA | 0.400 | 0.429 |
| VoroIF-GNN | 0.223 | 0.247 |
| TopoQA | 0.212 | 0.205 |
| GuijunLab-QA* | 0.138 | 0.151 |
| GuijunLab-Pathreader* | 0.148 | 0.142 |
| MULTICOM* | 0.139 | 0.136 |
| MULTICOM_GATE* | 0.139 | 0.132 |
| MULTICOM_LLM* | 0.139 | 0.128 |
| ModFOLDdock2* | 0.155 | 0.149 |
| MQA_base* | 0.150 | 0.144 |
| ContrastQA | **0.123** | **0.116** |

Note: Data in bold indicates the best result. Those with \* refer to the methods of the teams participating in CASP16, where the data are extracted from the official website results.

As shown, our method achieved outstanding performance in both TMscore and GDT-TS evaluations. Specifically, we obtained the lowest TMscore *Top1 loss* of 0.123, which is 0.015 (10.9%) lower than the best-performing method in CASP16 (GuijunLab-QA) with a ranking loss of 0.138, and 0.025 (16.9%) lower than the best-performing single-model method in CASP16 (GuijunLab-PAthreader) with a ranking loss of 0.148. The results are also comparable to most top-performing multi-model methods, demonstrating the excellence of our method’s performance. On GDT-TS *Top1 loss* we reached 0.116, achieving the lowest ranking loss among all 12 methods, which is also 8.7% lower than the second place MULTICOM LLM and 17.7% lower than GuijunLab-PAthreader. This shows the excellent performance of ContrastQA in terms of ranking loss.

In SI Appendix, Tables.S2–S3 show the specific TMscore as well as the GDT-TS ranking loss of all CASP16 methods on each target of the CASP16 dataset, respectively. In the TMscore section, it can be seen that ContrastQA achieved the lowest ranking loss for 9 targets among 37 targets. In particular, on the H1217, H1227, and H1272 large multimers reached 0.020, 0.001, and 0.053, respectively, which illustrates the excellent performance of ContrastQA in accurately assessing the model quality of large multimers. Therefore, to explore the superiority of ContrastQA for large multimers, we further conducted a case study of target H1217 on this, as shown in SI Appendix, FigS2. In the GDT-TS evaluation, ContrastQA achieved the lowest ranking loss on 8 out of 37 targets. Notably, the loss for target H1272 was 0, and for H1227 it was 0.062—significantly lower than those of other methods. These results highlight the particular advantage of our method in assessing large multimers.

To further explore the performance of our method on specific targets, we compared the performance of all methods on hard targets as well as easy targets based on the difficulty distinction of targets in CASP16. An easy target typically has available homologous templates, resulting in high-accuracy predictions, good model quality, and strong differentiation among decoys. In contrast, a hard target lacks homologous templates, leading to low-quality predictions, poor differentiation, and low similarity to the native structure, thus posing a greater challenge. Therefore, if the structure quality in hard targets can be predicted effectively and accurately, it can also further indicate that the EMA method has the potential for in-depth research on multimers without available homology templates, such as orphan proteins. We first compared the performance of all methods on hard targets in terms of TMscore *Top1 loss*, as shown in Table 2.

**Table 2.** TMscore ranking loss of hard targets on the CASP16 dataset.

| Target | ContrastQA | GuijunLab-QA* | GuijunLab-Pathreader* | MULTICOM_LLM* | ModFOLDdock2* | MQA_base* | TopoQA |
| --- | --- | --- | --- | --- | --- | --- | --- |
| T1201o | 0.011 | 0.013 | 0.027 | 0.014 | 0.010 | 0.007 | 0.009 |
| T1218o | 0.475 | 0.584 | 0.117 | 0.635 | 0.634 | 0.127 | 0.558 |
| T1237o | 0.009 | 0.011 | 0.009 | 0.007 | 0.008 | 0.008 | 0.014 |
| T1240o | 0.379 | 0.326 | 0.655 | 0.357 | 0.236 | 0.386 | 0.389 |
| T1249o | 0.020 | 0.151 | 0.027 | 0.082 | 0.117 | 0.025 | 0.752 |
| T1257o | 0.018 | 0.005 | 0.005 | 0.017 | 0.015 | 0.731 | 0.731 |
| T1269o | 0.047 | 0.027 | 0.027 | 0 | 0.041 | 0.047 | 0.029 |
| T1270o | 0.213 | 0.210 | 0.227 | 0.206 | 0.136 | 0.218 | 0.211 |
| T1295o | 0.119 | 0.119 | 0.119 | 0.259 | 0.263 | - | 0.256 |
| T1298o | 0.019 | 0.297 | 0.327 | 0.295 | 0.310 | 0.009 | 0.312 |
| Average | **0.131** | 0.174 | 0.154 | 0.187 | 0.177 | 0.173 | 0.326 |

Note: Data in bold indicates the best result. Those with \* refer to the methods of the teams participating in CASP16, where the data are extracted from the official website results. “-” means since MQA\_base did not evaluate T1295o, we only counted the remaining nine results from MQA\_base.

Table 2 shows that ContrastQA can estimate model quality accurately well for hard targets, achieving the lowest average ranking loss: 0.131, which is 0.023 (14.9%) and 0.042 (24.3%) lower than the second and the third best performing method-GuijunLab-PAthreader and MQA base-with ranking losses of 0.154 and 0.173. As can be seen, we achieved a more balanced performance as the output suggests. To further explore our strength, we first checked the TMscore distributions of all hard targets, and eventually T1249o caught our attention, with a polarized distribution, i.e., many are low-quality structures and some are high-quality structures, as shown in Fig. 2A. Therefore, we selected T1249o from hard targets for in-depth experimentation because we achieved the lowest TMscore ranking loss of T1249o among all CASP16 EMA methods. In Fig. 2B, the three protein multimers shown from left to right are: the native structure, the top-ranked model based on the true TM-score, and the top-ranked model selected by ContrastQA, respectively. We can see that the model chosen by ContrastQA reaches a good similarity with the native structure. This model has a high TMscore of 0.9591 and the TMscore ranking loss is much lower than CASP16 EMA methods. This means that our method has great potential in exploring the field of assessing the quality of template-free proteins. We also compared the performance of the methods on easy targets, as shown in the SI Appendix, Table.S4, and it can be seen that ContrastQA still performed well.

**Fig. 2.**
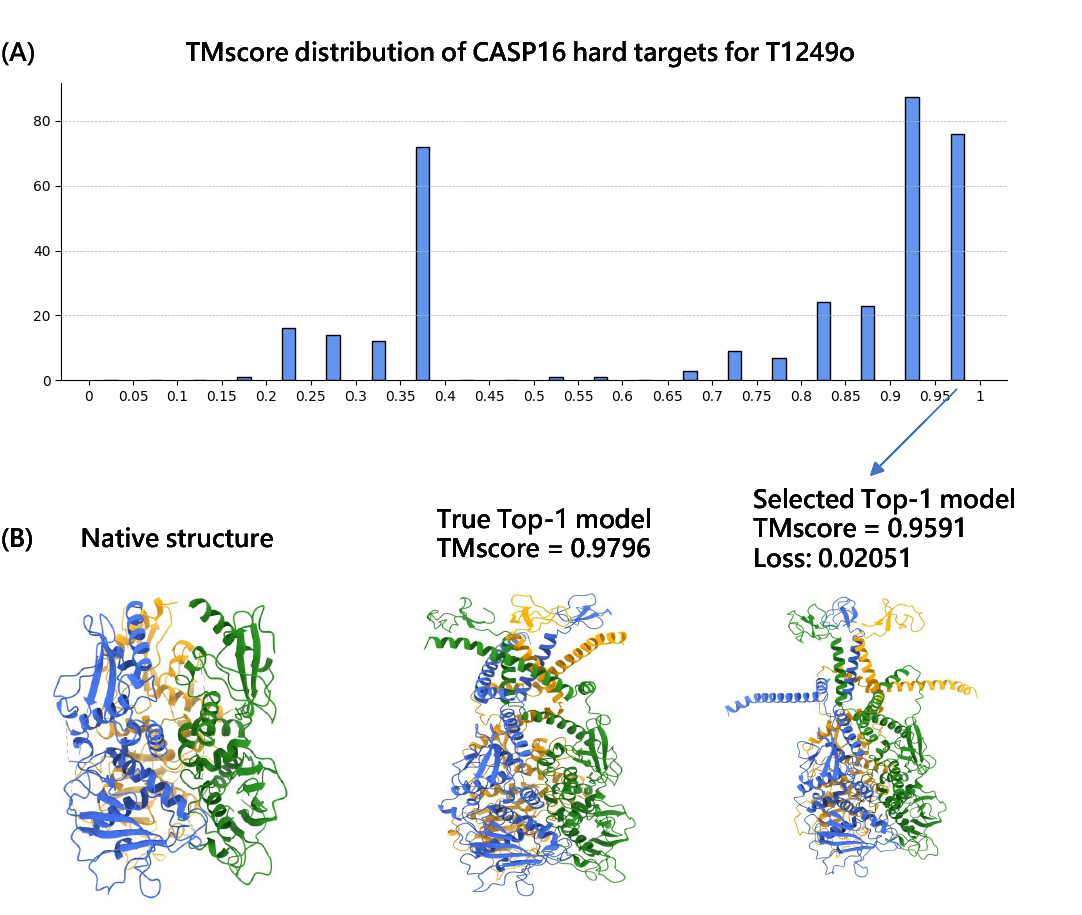
**(A)** is the TMscore distribution of CASP16 hard targets for T1249o. **(B)** includes corresponding native structure, the true TOP-1 model (TMscore is 0.9796) and the ContrastQA Top-1 selected model (TMscore is 0.9591, ContrastQA achieved a 0.02051 TMscore ranking loss.) to target T1249o.

### Performance on the ABAG-AF3 dataset

Following the approach TopoQA, ABAG-AF3 is based on the latest protein structure prediction method, Alphafold3 (1), which set five different seeds to run five times to derive the decoys based on the input protein sequence. The predicted structures have high quality and low discriminatory. We tested ContrastQA on 35 targets on the ABAG-AF3 dataset and compared it with multiple deep learning-based methods for EMA of protein complexes.

As shown in Table 3, ContrastQA achieved leading results in terms of ranking loss. Our ranking loss on TMscore is 0.028, which is 9.7% lower than that of VoroIF-GNN with a ranking loss of 0.031, and 24.3% lower than that of the third place TopoQA with a ranking loss of 0.037. For GDT-TS, our ranking loss is 0.021, which is 19.2% lower than TopoQA with a ranking loss of 0.026, and 22.2% lower than VoroIF-GNN with a ranking loss of 0.027. It can be seen that the excellent performance of our method on the AF3 dataset. This coincides with the conclusion we reached in the previous section: namely, the salient advantages of our EMA on the structures predicted by the novel protein multimer structure prediction methods.

**Table 3.**
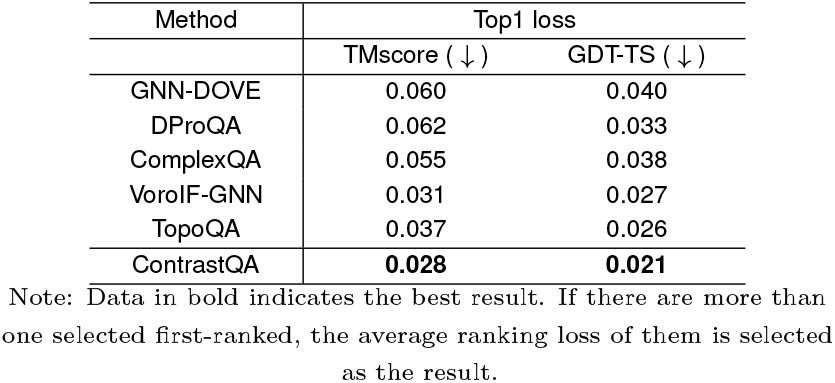
Performance on the ABAG-AF3 dataset.

In SI Appendix, Tables.S6–S7 specifically show the specific performance of ContrastQA with the five SOTA methods on the TMscore as well as the GDT-TS ranking loss. In the TMscore section, a much lower ranking loss was achieved on the 8 targets, especially on 7su0, 7su1 as well as 7x7o which achieved the ranking loss of 0, indicating the successful selection of the correct first-ranked structure. On 7wrv, our method achieved a ranking loss of 0.062, which is much lower than the other methods. In terms of GDT-TS, our method achieved the lowest ranking loss on 9 out of 35 targets.

Notably, it achieved a loss of 0 on targets 7r40, 7su1, and 7yqx. Even on challenging targets such as 7wrv, it attained the lowest loss of 0.003, which is significantly lower than those of other methods. Generally, several metrics show the outstanding potential of ContrastQA in evaluating protein complex structures predicted by novel high-quality protein prediction tools, which can find out the better one among many conformations under the same target.

### Ablation study

In order to evaluate the impact of the label-guided graph contrastive learning module and the specially designed contrast loss function on our method, we performed ablation experiments by removing relevant specific sections, and the results are shown in Table 4. W/o gcl module denotes the removal of the graph contrastive learning module and the training of the model using only the geometric graph neural network. W/o weighted cl loss indicates the modification of the contrast loss function part, removing the weights before the negative samples in the computation process, uniformly all set to 1.

**Table 4.** Ablation study on CASP16 dataset and ABAG-AF3 dataset.

| Dataset | Method | Top1 loss (↓) |  |
| --- | --- | --- | --- |
|  |  | TMscore | GDT-TS |
| CASP16 | w/o gcl | 0.198 | 0.170 |
|  | w/o weighted cl loss | 0.186 | 0.189 |
|  | ContrastQA | **0.123** | **0.116** |
| ABAG-AF3 | w/o gcl | 0.039 | 0.027 |
|  | w/o weighted cl loss | 0.031 | 0.024 |
|  | ContrastQA | **0.028** | **0.021** |

Note: Data in bold indicates the best result.

We first removed the graph contrastive learning module and trained the model using only the geometric graph neural network GVP-GNN. The results show that the performance of the method is significantly degraded. The TMscore ranking loss worsened by 0.075 on the CASP16 dataset, increasing by 37.9%, and the GDT-TS ranking loss elevated by 31.8%. On the ABAG-AF3 dataset the TMscore ranking loss increased by 28.2% from 0.028 to 0.039 and the GDT-TS ranking loss was elevated by 22.2%.

We then validated our designed contrast loss function by removing the weights before the negative samples from the calculation process, ignoring the difference of whether it is the same target or not, and uniformly setting them all to 1. From the results, we can see the impact of the designed loss function: the TMscore ranking loss on the CASP16 dataset increased from 0.123 to 0.186, a deterioration of 0.063 (33.9%), and the GDT-TS ranking loss (0.116) achieved 0.073 (38.6%) lower than w/o weighted cl loss with a ranking loss of 0.189. On the ABAG-AF3 dataset, the TMscore and GDT-TS ranking losses increased from 0.028 to 0.031 and from 0.021 to 0.024, representing deterioration of 0.003 (9.7%) and 0.003 (12.5%), respectively.

We further investigated the role of contrastive learning by examining the impact of each key parameter, as illustrated in Fig. 3. We began with the temperature coefficient, considering the limited variability between anchor and negative samples in our data. Using 0.5 as the initial value, we observed a significant performance drop—TMscore ranking loss increased by 0.09 (42.3%) compared to the optimal value of 0.123. This decline is attributed to the overly high tolerance for negative samples, which impaired the model’s ability to effectively distinguish hard negatives. As the temperature coefficient gradually decreased, we observed a corresponding decrease in the ranking loss. This suggests that paying more attention to hard samples in our task helps better distinguish between samples with subtle differences. The main reason behind this effect is that most of the protein structures in our dataset are of high quality and exhibit very little variation from one another.

**Fig. 3.**
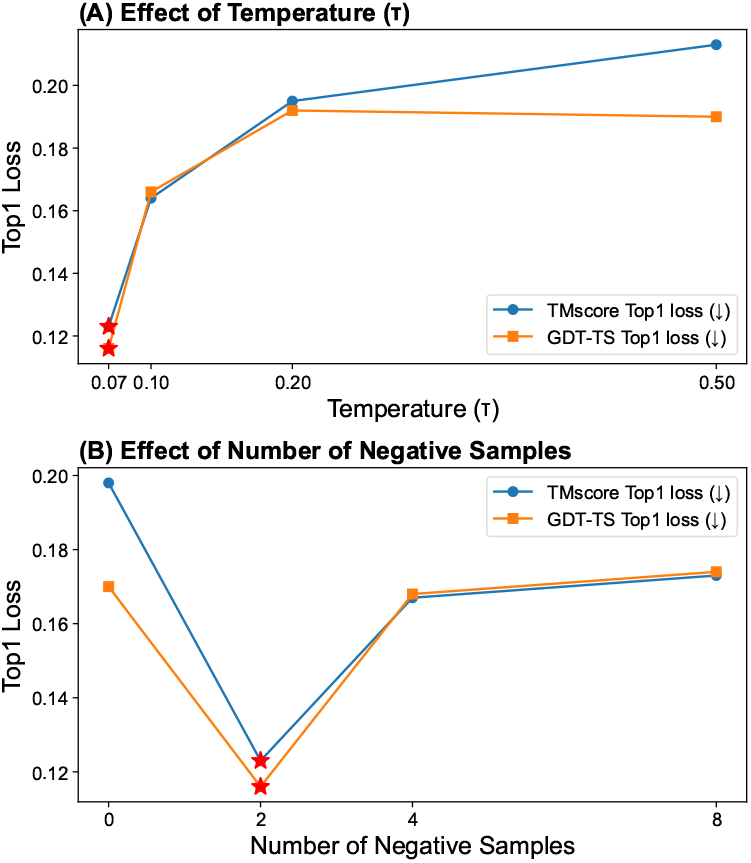
Fine-grained ablation study of ContrastQA. **(A)** illustrates the variation of ContrastQA’s performance in terms of temperature coefficient, with the effect deteriorating as the temperature coefficient continues to increase. **(B)** demonstrates the change in the effectiveness of ContrastQA with respect to the number of negative samples, and it can be seen that the contrastive learning greatly improves the effectiveness (number 2 vs. number 0). After number of 2, the effectiveness is greatly reduced as the number of negative samples continues to increase. All other parameter settings are consistent with the original experimental parameter settings in this paper. In particular, due to hardware limitations, we cannot further increase the number of negative samples beyond 8.

On the other hand, we also explored the effect of the number of negative samples, due to the limitation of the hardware, we can only set the number of negative samples to a maximum of 8. We know that when the number of negative samples gradually increases, the effect of contrastive learning should be improved, because the distance between the anchor sample and the negative samples can be widened. However, since the vast majority of protein structures in our dataset are of high quality, selecting negative samples becomes more challenging. When the number of required negative samples increases (where negative samples are categorized as Incorrect), more of them need to be drawn from other targets. This leads to the model placing excessive focus on the differences between the anchor sample and negative samples from different targets. However, structural differences across different targets are already large by nature. As a result, the model pays less attention to the subtle differences between the anchor sample and negative samples within the same target, which are more relevant for fine-grained discrimination. The situation may be entirely different when the negative sample belongs to the correct category. However, as this scenario is relatively rare, our experimental design primarily focuses on the more frequent case where negative samples are incorrect. The main task of protein EMA is still to select the best quality decoy from the same target, so we should try to pay more attention to the difference between the anchor sample and the negative samples under the same target and increase the distance between the two. The experimental results also confirmed our conjecture, i.e., it is not the case that more negative samples enhance the effectiveness of our task. The ranking loss with a negative samples size of 2 is 0.05 (28.9%) and 0.058 (33.3%) lower than the ranking loss with a size of 8. Also we can find that the ranking loss worsened by 0.075 (37.9%) and 0.054 (31.8%) when the number of negative samples is 0, i.e., when contrastive learning is not used.

In summary, both the graph contrastive learning module as well as the designed contrast loss function have a significant impact on the model performance. Among them, it can be seen that the graph contrastive learning module has greater impact, with the sum of ranking loss increased up to 35.1% as well as 25.8% on CASP16 and ABAG-AF3 datasets, respectively. It is enough to verify the powerful protein graph representation learning ability of the designed loss function and the proposed graph contrastive learning module.

## Discussion

In this study, we proposed ContrastQA, a novel protein model EMA method utilizing a novel graph contrastive learning framework combined with geometric graph neural networks. The division of positive and negative samples was guided by using the interface score, DockQ, as a score label, and was also used as an important component of contrast loss to guide graph contrastive learning during subsequent model training. The ablation study indicated that our proposed graph contrastive learning framework significantly improved model testing performance. After removing the graph contrastive learning module and changing the contrast loss function, the performance of our method decreased compared to the original method on two testing sets respectively.

In the evaluation of global quality for predicted protein multimers, our method demonstrated strong competitiveness across multiple datasets. On the latest CASP16 dataset, it outperformed a range of advanced EMA methods, achieving the lowest TMscore ranking loss of 0.123 and GDT-TS ranking loss of 0.116—representing improvements of 10.9% and 8.7% over the second-best methods, GuijunLab-QA and MULTICOM LLM, respectively. Furthermore, on the AF3 dataset, which features low structural variability and high global quality, our method continued to perform well, with TMscore and GDT-TS ranking losses 9.7% and 19.2% lower than the second-best results. It can be demonstrated the feasibility of our proposed graph contrastive learning module in the field of protein model EMA.

It is worth noting that our method was trained and validated on a relatively small dataset comprising approximately 20,000 decoys, which is significantly smaller than the datasets used by CASP16 teams—for example, GuijunLab-QA was trained on nearly 2 million decoys. We know that graph contrastive learning produces better generalization on larger datasets, and we can foresee that the potential of ContrastQA can be exploited to a greater extent in the future if the dataset size can be expanded.

The graph contrastive learning module in ContrastQA currently defines positive-negative samples pairs based solely on the interface score DockQ. However, relying on DockQ alone may lead to a somewhat biased partitioning. In future work, we aim to incorporate physicochemical interface features alongside DockQ to achieve a more balanced and robust sampling strategy. Meanwhile, our method mainly focused on the global accuracy estimation, and did not involve the interface and local EMA. In the future, we hope to extend the assessment field to protein multimer interface and even local quality evaluation by combining with more fine-grained multi-task learning.

## Materials and Methods

### Datasets

The SI Appendix section of Distribution of training, testing, and validation datasets and Table 5 show the specific distribution of the training, validation, and testing datasets, with each target associated with multiple decoys. Among them, we computed datasets’ score labels using DockQv2 (24).

**Table 5.** Distribution of training, testing, and validation datasets.

| | Dataset | Num targets | Num decoys | Correct(DockQ ≥ 0.23 ) | Incorrect(DockQ < 0.23 ) |
| --- | --- | --- | --- | --- | --- |
| Training | MULTICOM_train | 32 | 3200 | 89.87% | 10.13% |
| Training | PPI-AF2_train | 56 | 1400 | 89.93% | 10.07% |
| Training | PPI-AF3_train | 354 | 7080 | 94.48% | 5.52% |
| Training | Voro_train | 678 | 8102 | 41.43% | 58.57% |
| Validation | MULTICOM_val | 8 | 800 | 87.50% | 12.50% |
| Validation | PPI-AF2_val | 14 | 350 | 96.00% | 4.00% |
| Validation | PPI-AF3_val | 89 | 1780 | 89.33% | 10.67% |
| Validation | Voro_val | 166 | 2024 | 42.54% | 57.46% |
| Testing | CASP16 | 37 | 12142 | 81.04% | 18.96% |
| Testing | ABAG-AF3 | 35 | 875 | 93.49% | 6.51% |

This table shows the distribution of the training validation as well as the testing set. The targets denote protein structures with independent amino acid sequences, each target has multiple decoys, i.e., these decoys have the same amino acid sequence but different 3D structures. Each decoys will be labeled with its own score in preparation for subsequent tasks.

#### Training and validation datasets

To train the ContrastQA model, we first collected MULTICOM Dataset which is a protein quaternary structure dataset constructed by MULTICOM (29) generation. We then filtered 40 targets where each target contains 100 decoys, leading a total of 4,000 decoys based on the level of quality reflected in the DockQ (24) score labeling. Then we constructed PPI Dataset based on the amino acid sequences in PPI4DOCK (30) using AF2-Multimer (31) as well as AF3 (1), which generated 70 targets totaling 1,750 decoys by using AF2-Multimer, and AF3 generated 443 targets totaling 8,860 decoys. Additionally, we incorporated 10,126 decoys for 844 heterodimeric targets from VoroIF-GNN (14). Subsequently, we trained our model by combining the three datasets, **MULTICOM Dataset, PPI Dataset, Voro Dataset**. We used MMseq2 (32) to compare the sequence similarity of the MULTICOM, PPI, and Voro datasets with the native structures, and filtered out sequences containing residues inconsistent with the native structures. To ensure low sequence redundancy, we selected training and validation sets such that the sequence similarity between them is below 30%, with a roughly 8:2 split ratio.

#### Testing datasets

To benchmark the performance of our method ContrastQA with many SOTA methods, we collected 12,142 decoys for 37 protein multimer targets from CASP16, which is **CASP16 Dataset**. We also employed the **ABAG-AF3 dataset** introduced in TopoQA (16), which contains structures predicted by the SOTA protein complex prediction method AlphaFold3. Most of these structures are of high quality, with minimal structural deviations, and 93.5% of them are classified as Correct (DockQ *≥* 0.23). The ABAG-AF3 dataset contains 35 protein multimer targets with 875 decoys. To show the stringency, the sequence similarity between the training validation sets and the testing set is set to less than 25%.

### A Label-guided Graph Contrastive Learning-based Framework

The flowchart of our method is shown in Fig. 1A. It consists of four main parts: data preprocessing, positive-negative samples pairs division, graph contrastive representation learning module, and global quality score prediction. We first extracted features from the input complex structure in terms of both sequence as well as structure, including ESM-2 sequence embedding features (33), DSSP features (34), triangle position features (15), sequence one-hot features, laplacian positional encoding features (35), edge position encoding features (36, 37), chain ID features, contact features, and atomic pairwise distance features (13). After feature fusion, we utilized DGL (38) to save as a dgl graph data structure. Each input structure was divided into positive-negative samples pairs based on DockQ (24) scores by our designed positive-negative samples pair selection strategy. In order to improve the learning effect of protein structure representation, we introduced the designed graph representation contrastive learning module. All samples obtained in the previous section were fed into GVP-GNN (23) to obtain protein graph embeddings. We used global average pooling in DGL to aggregate node features into graph-level representations. Each input structure, along with its corresponding positive and negative samples, were then used for contrastive learning. Finally, the input structure outputted its corresponding global quality score through a multilayer perceptron (MLP).

#### Positive-negative samples pair selection strategy

The traditional contrastive learning methods (as illustrated in Fig. 1B) mostly use data augmentation, perturbation and other means to generate positive and negative samples (19, 20), and often divide them randomly without considering whether they originate from the same target or share similar scores label. Given the specificity of the protein EMA task, we argued that it is essential to incorporate both the target identity and the score label as key criteria for constructing positive and negative sample pairs. The former enables fine-grained differentiation within the same protein system, while the latter provides precise guidance based on structural quality. We designed a novel positive-negative samples pair selection strategy, as shown in Fig. 1C, for which we labeled the scores based on the interface quality score, DockQ (22, 24), and measured the quality of protein complexes in the range of [0,1], with larger values indicating better accuracy. Since our decoys involved complexes of trimers and even more chains, we used the latest DockQv2 (24), which can score multiple interfaces as a whole, as our score labels.

It is specified that DockQ has four categories as follows: (0,0.23) is Incorrect, [0.23,0.49) is Acceptable quality, [0.49,0.8) is Medium quality, and [0.8,1] is High quality. We further divided it into two categories, Incorrect vs. Correct (Acceptable, Medium and High quality). There are multiple decoys per target in our datasets. For each input sample, we first determined its score category (Correct or Incorrect). If the sample is categorized as Incorrect, the positive sample was randomly selected from other Incorrect decoys of the same target. Negative samples were first drawn from Correct decoys of the same target; if insufficient, additional Correct decoys were randomly selected from different targets. Conversely, if the input sample was in the Correct category, the same strategy applied in reverse. We specify that each group consists of an anchor sample, one corresponding positive sample, and multiple corresponding negative samples.

#### Graph contrastive learning module

Graph contrastive learning is to bring similar graphs closer to each other and different graphs farther away from each other, so as to effectively learn the features and structures in graph data (18, 19, 39). Most traditional graph contrastive learning methods are based on unsupervised learning. In our work, we modified this approach to better suit the specificity of our task. Specifically, we utilized DockQ score labels to guide the construction of positive-negative samples pairs. Based on these labels, we normalized the data and then applied graph contrastive learning to the constructed sample pairs.

As shown in Fig. 1A, based on the positive-negative samples pair selection strategy mentioned in the previous section, we designed a label-guided graph contrastive learning module. For the KNN graph after feature extraction of the input structure, the positive-negative samples pair belonging to it was classified according to the DockQ score labels, and each of them was subsequently fed into the corresponding geometric graph neural network to learn the graph embedding representation. In order to better represent the three-dimensional structure of proteins and at the same time cope with the rotational invariance of proteins, we introduced a geometric graph neural network thus being able to learn protein feature efficiently. GVP-GNN (23) adds feature directionality to GNN, enabling the model to optimize scalar features using vector features, better capture complex representations, and more accurately describe protein 3D structures. The node and edge features were fed into the GVP to compute and pass the information respectively, and the final graph representation was used as input to the GVP-GNN for subsequent message passing. After that, GVP-GNN computed scalar and vector features together and integrated the orientation information into the scalar features, finally, we applied a layer of GVP for refining the feature representation of proteins, which is helpful to improve the model performance. The final was to get the scalar features of the nodes as graph embedding representations.

After obtaining the graph embedding representation, the final graph representation was obtained for each sample through graph global average pooling strategy. At this point, the graph representations of the input structure and its corresponding positive-negative samples were proposed through graph contrastive learning to obtain the contrast loss. For the purpose of maximizing the effect of graph contrastive learning, we constructed the NT-Xent loss (39) function based on the traditional NT-Xent loss function applicable to our own task, in which the positive-negative samples pairs are explicit inputs.

Specifically, for a given target, high-quality decoys tend to share similar 3D structures, while low-quality decoys often exhibit distorted or implausible conformations, which aligns with the core idea of contrastive learning (7). Importantly, when negative samples are drawn from different targets, the inherent sequence dissimilarity ensures that these samples are generally less similar to the anchor sample than negatives selected from the same target. Therefore, it is critical to distinguish between these two cases. By leveraging the DockQ score labels, we can incorporate the score difference into the loss function. Specifically, the difference in DockQ scores between the input sample and its corresponding negative samples can be used as a weighting factor for the negative samples in the loss computation. Given the idea of multi-granularity, different treatments were given in the scenario of whether it is the same target or not. If the negative samples belong to the same target as the input sample, we set the difference between the DockQ scores as the weight of the negative samples before comparing the loss, and if the situation is reversed, we set the weight of the negative samples to 1. Therefore, it can reflect as much as possible the differences between the negative samples under the same target and different targets.

The specific loss function is expressed as formula 1:

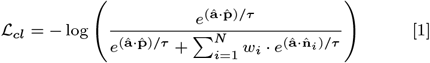

where 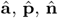 denote the anchor sample, the positive sample, and the negative sample, respectively and 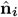 denotes the i^th^ negative sample. The *τ* (i.e., temperature) (18) is used in contrastive learning to control the emphasis placed on hard negative samples. A larger temperature value results in a smoother contrastive loss, which reduces the focus on difficult negative samples. In contrast, a smaller temperature value increases the model’s attention to hard negatives — those that are highly similar to the anchor sample. In this case, the model becomes more “serious” in distinguishing between positive and negative samples. The weight of the loss of contrast between the anchor sample and the i^th^ negative sample is denoted as *w*_*i*_ which has the following formula 2:

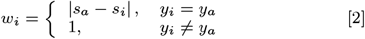

where *s*_*a*_, *s*_*i*_ represent the DockQ scores of the anchor sample as well as the i^th^ negative sample, respectively. Meanwhile, the *y*_*i*_ and *y*_*a*_ indicate whether the i^th^ negative sample and the anchor sample belong to the same target.

### Feature design

As shown in Fig. 1A, the protein complex structure is encoded as a KNN (10-neighbor) graph, whose nodes denote amino acid residues and edges between nodes denote a residue pointing to its K nearest neighboring residues in the 3D structure (based on the distance of C_*α*_ atoms), and the model graph contains node features and edge features. The node features have a total of 512 dimensions and are used to characterize the protein residue information. Edge features with a total of 22 dimensions are used to characterize the interactions between protein residues.

#### Node features

In order to effectively characterize protein residues, the node features consist of two parts: protein base features and high-dimensional features generated by Protein Language Model (PLM). The base features are 55 dimensions in total, including one-hot encoding features of secondary structure, backbone torsion angle (PHI and PSI) features using sine and cosine values, and solvent accessibility surface area features of 14 dimensions, which were generated by DSSP program (34). We used a 21-dimensional one-hot encoding to represent amino acid sequences. Following DProQA (13), we also incorporated an 8-dimensional Laplacian positional encoding to capture the spatial positioning of residues. In addition, to better represent the relationship between local and global topological features of residues, we adopted the 12-dimensional triangular position encoding proposed by GraphCPLMQA (15).

Additionally, protein language models (PLMs) (12, 33, 40) capture evolutionary patterns from large-scale protein sequence datasets. These models are capable of representing potential relationships between sequences in high-dimensional spaces and have been widely applied in protein structure prediction and functional analysis. Therefore, PLMs have been widely adopted to generate robust embeddings for protein structural feature extraction (33, 40, 41). We adopted ESM-2 (33) to extract amino acid sequence embedding features, and extracted the 1280-dimensional features in the last layer of the network (Layer 33) as the residue-level embedding features. In order to well fuse the base features as well as the ESM features, we input both to the MLP mapping to the same feature dimension and performed feature fusion, finally obtained 512-dimensional node features.

To capture the three-dimensional structural information of the protein—specifically the relative positions and orientations of individual residues—we computed a 2D orientation feature of the backbone. This vector-based feature was derived from the 3D coordinates of the residues’ C_*α*_ atoms (42).

#### Edge features

The edges in the KNN graph reflect the relationship between a residue and its neighboring neighbors in terms of C_*α*_ atomic distances. The edge features are 22 dimensions in total. First we introduced a 1-dimensional contact feature to define the contact between residues (i.e. if C_*β*_ - C_*β*_ atomic distance *≤* 8*Å*). Then we introduced 3 distance features (i.e. C_*α*_-C_*α*_ atomic distance, C_*β*_ - C_*β*_ atomic distance, and the distance between backbone nitrogen and oxygen atoms of two residues). Finally, the edgewise positional encoding (36) and relative positional encoding (37) were represented, they together added up to the positional encoding feature of 17 dimensions.

### Model training

We trained model using a combination of structure global quality loss and graph contrastive learning loss terms. Specifically, in the structure global quality loss, based on the score label lDDT (43), we referred to the category division of pLDDT in AlphaFold2 (3) to divide similar categories for lDDT, i.e., lDDT < 0.5 is regarded as VERY LOW, 0.5 ⩽ lDDT < 0.7 is regarded as LOW, and 0.7 ⩽ lDDT < 0.9 is regarded as CONFIDENT, 0.9 ⩽ lDDT is VERY HIGH. Thus, there are two global quality losses, the cross-entropy loss with respect to category adoption, where the loss term is quadratic, and the mean square error loss (MSE), which was used to compute the square loss between the predicted global quality scores and the reference global quality scores. For the graph contrastive learning loss, we used the modified NT-Xent loss (39). The total loss is then the sum of the global quality loss and the graph contrastive learning loss as follows:

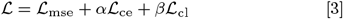

where *ℒ* corresponds to the individual loss functions, the coefficient *α* represents the weight of the quadratic cross-entropy loss, and the coefficient *β* represents the weight of the graph contrastive learning loss, both of which are constants. In this paper, we set *α* = 0.1 and *β* = 0.005. In SI Appendix section of Training Details and Loss Learning Curve and Table.S1, Figs.S1(A)-(D) show how we trained, tuned ContrastQA and the specific training process.

We used the Adam (44) optimizer as the optimizer with a learning rate of 1e-3 and a weight decay rate of 5e-4. We trained the model on an NVIDIA RTX 4090D GPU (24GB video memory) device for 50 epochs with the batch size set to 16. To mitigate the overfitting we also set up an early-stopping strategy, where we stopped the training early when the loss on the validation set did not decrease within 10 consecutive generations.

## Data, Materials, and Software Availability

The data of CASP16 and ABAG-AF3 datasets are available at https://predictioncenter.org/download_area/ and http://mialab.ruc.edu.cn/ABAG-AF3/zip, respectively. The data of training dataset and pre-trained model are available at https://dx.doi.org/10.5281/zenodo.15463235.

## ACKNOWLEDGMENTS

This work was supported by the Natural Science Foundation of Anhui Province (No. 2408085MF152) and the Key Projects of the University Excellent Talents Support Plan of Anhui Provincial Department of Education (No. gxyqZD2021089).

