## Supplemental file for "ContrastQA: A label-guided graph contrastive learning-based approach for protein complex structure quality assessment"

**This PDF file includes:**

Figs. S1 to S2
Tables S1 to S7
SI References

### Training details and loss learning curve.

We trained the ContrastQA model using the PyTorch(1), PyTorch Lightning(2) framework with multi-task learning, and the training loss consists of three components: the Mean Squared Error (MSE) loss between the predicted scores and the true scores, the Cross-Entropy (CE) loss between the predicted score categories and the true score categories, and the contrast loss. We chose the best model based on the validation set's performance. All parameters' settings are described in Table S1.

The formula for MSE loss is as follows:

$$\mathrm{MSE} = \frac{1}{N}\sum_{i=1}^{N}(y_i - \hat{y}_i)^2 \qquad [1]$$

where N is batch size, $y_i$ is the ture score of the $\mathrm{i^{th}}$ sample, $\hat{y}_i$ is the predicted score of the $\mathrm{i^{th}}$ sample, $(y_i - \hat{y}_i)^2$ is the squared error between the predicted and true values of the $\mathrm{i^{th}}$ sample.

The formula for CE loss is as follows:

$$\mathrm{CE} = -\frac{1}{N}\sum_{i=1}^{N}\log\left(\frac{e^{z_{t_i}^{(i)}}}{\sum_{k=0}^{3} e^{z_k^{(i)}}}\right) \qquad [2]$$

where N is the batch size, $z_k^{(i)}$ is logit of the $\mathrm{i^{th}}$ sample for the $\mathrm{k^{th}}$ class, $t_i \in [0,1,2,3]$ is the index of true categories.

The learning curves of the three losses during model training are represented in Fig.S1(A)-(D).

### Distribution of training, testing, and validation datasets

We refer to the datasets provided by MULTICOM(3), PPI4DOCK(4), and VoroIF-GNN(5), and obtain our dataset after subsequent screening for sequence similarity, quality issues, and length issues. When the sequence similarity is below 30% by MMseq2(6), a quality issue refers to cases where the decoy has a significantly different amino acid sequence from the reference and shows inconsistent labeling between the DockQ(7) and lDDT(8) scores. A length issue refers to the exclusion of decoys with more than 1000 amino acid residues. The test dataset includes CASP16 as well as ABAG-AF3(9), and we set the sequence similarity threshold to 25% between it and the training validation set. For the decoys with sequence lengths greater than 1000 amino acid residues, we processed them individually to generate ESM-2(10) embedded features, and we deleted the decoys that could not be generated into intermediate feature files.

### Evaluation metrics

We adopted TMscore (11) as well as GDT-TS (12) as evaluation metrics, both of which are used to assess the global quality of the protein structure. We use US-align(13) to compute the TMscore and openstructure(14) to compute the GDT-TS.

The GDT-TS score is defined as the average percentage of $\mathrm{C}_\alpha$ atoms in the predicted structure that fall within a certain distance cutoff from their corresponding atoms in the native structure, under four progressively increasing thresholds: $1\AA$, $2\AA$, $4\AA$, and $8\AA$. Formally, the GDT-TS score is calculated as:

$$\mathrm{GDT-TS} = \frac{1}{4}\left(\mathrm{GDT}_1 + \mathrm{GDT}_2 + \mathrm{GDT}_4 + \mathrm{GDT}_8\right) \qquad [3]$$

where $\mathrm{GDT_n}$ denotes the percentage of residues in the predicted model whose $\mathrm{C}_\alpha$ atoms are within $\mathrm{n}\AA$ of the corresponding residues in the reference structure. The GDT-TS score ranges from 0 to 100, with higher values indicating greater structural similarity. In the article we mapped it to [0,1] for ease of computation. Unlike RMSD, GDT-TS is more robust to local deviations and partial alignments, making it a more reliable indicator for assessing global model quality.

TMscore (Template Modeling score) is a widely adopted metric to assess the topological similarity between two protein structures. Unlike RMSD and GDT-TS, TMscore is more sensitive to global structural similarity and less affected by local errors. It ranges between (0, 1], with higher values indicating greater structural resemblance. The TMscore is computed as:

$$\mathrm{TMscore} = \max\left[\frac{1}{L_{\mathrm{ref}}}\sum_{i=1}^{L_{\mathrm{ali}}}\frac{1}{1+\left(\frac{d_i}{d_0}\right)^2}\right] \qquad [4]$$

where $L_{\mathrm{ref}}$ is the length of the reference structure, $L_{\mathrm{ali}}$ is the number of aligned residue pairs, $d_{\mathrm{i}}$ is the distance between the $\mathrm{C}_\alpha$ atoms of the $i^{\mathrm{th}}$ aligned residue pair, $d_0$ is a scale to normalize the match difference. By design, TMscore is normalized to avoid length bias and gives a score of 1 for a perfect match. Empirically, TMscore >0.5 suggests the two structures share the same fold.

### Testing specific results

**The specific TMscore as well as the GDT-TS ranking loss of CASP16 methods on each target of the CASP16 dataset.** We compared with other state-of-the-art (SOTA) methods in the field on the 37 targets above CASP16, Table S2 and Table S3 show the specific performance of all the methods in terms of TMscore ranking loss as well as GDT-TS ranking loss, where smaller values indicate better predictions. The specific data of each target was intercepted from the CASP official website. Since we did not have access to the native structure for H1229 and H1230, we additionally tested H1265, T1219o, and T1295o.

For the CASP16 teams, we calculated the TMscore as well as the GDT-TS scores ourselves, using the rankings in their own target results, and included the results in the statistical comparison. In the TMscore section, it can be seen that ContrastQA achieved the lowest ranking loss for 9 targets among 37 targets. In the GDT-TS evaluation, ContrastQA achieved the lowest ranking loss on 8 out of 37 targets. Notably, the loss for target H1272 was 0, and for H1227 it was 0.062—significantly lower than those of other methods. These results highlight the particular advantage of our method in assessing large multimers.

**TMscore ranking loss of methods on easy target of the CASP16 dataset.** We also compared with other methods in the easy targets divided in CASP16, where the data of CASP teams were obtained from the official website. The specific results are shown in Table S4. It can be seen that ContrastQA still performed well.

**TMscore and GDT-TS top3 mean and top5 mean of all methods on each target of the CASP16 dataset.** We used Top3 and Top5 mean as an additional evaluation metric, Top3 and Top5 mean are the average of the true scores of the top three and five predicted decoy structures, respectively. The larger the value indicates the higher quality of the decoy structure and the better the method is.

We compared the Top 3 mean and Top 5 mean on the TMscore as well as the GDT-TS, and Table S5 reports the effect of all comparison methods on these two metrics. It can be seen that ContrastQA achieved the highest average scores on all four metrics, including a notable 0.737 on the Top3 mean of TMscore, which is 3.9% higher than the second-place DProQA. It reached a outstanding 0.609 on the Top3 mean of GDT-TS, 9.1% higher than the second-place TopoQA and 9.5% higher than the third-place VoroIF-GNN.

**The specific TMscore as well as the GDT-TS ranking loss of all methods on each target of the ABAG-AF3 dataset.** We compared with other advanced methods in the field on the 35 targets above ABAG-AF3 dataset, and Table S6 and Table S7 show the specific performance of all comparison methods in terms of TMscore ranking loss as well as GDT-TS ranking loss, where smaller values indicate better predictions. In the TMscore section, a much lower ranking loss was achieved on the 8 targets, especially on 7su0, 7su1 as well as 7x7o which achieved the ranking loss of 0, indicating the successful selection of the correct first-ranked structure. In terms of GDT-TS, our method achieved the lowest ranking loss on 9 out of 35 targets.

**Case study of ContrastQA on H1227 target.** As shown in Table S2 and Table S3, ContrastQA achieved the best results for TMscore as well as GDT-TS top1 loss of 0.001 and 0.062 at the target H1227 of CASP16, respectively. Target H1227 is a Hetero7-mer with a sequence length of 5689. As shown in Fig.S2A, it can be seen that the distribution of TMscore scores for H1227 is relatively concentrated, mostly in the high quality section (i.e., above 0.95). We therefore conducted a case study for this target, where we compared the model with the highest ContrastQA predicted scores with the model which has the highest actual TMscore scores, the native structure of H1227, all of them in three dimensions structures. The TMscore of the structure predicted by ContrastQA is 0.9839, while the top1 ranked TMscore is 0.9852. As shown in Fig.S2B, it can be seen that the ContrastQA Top-1 selected model is very similar to the native structure, even more so than the true Top-1 model. Specifically, the relative positions of the chains of the structure predicted by the ContrastQA remained consistent with the native structure, while the relative positions of the chains of the true Top-1 model did not coincide with the native structure. These results demonstrate the ability of ContrastQA to assess the quality of large multimers with high precision.

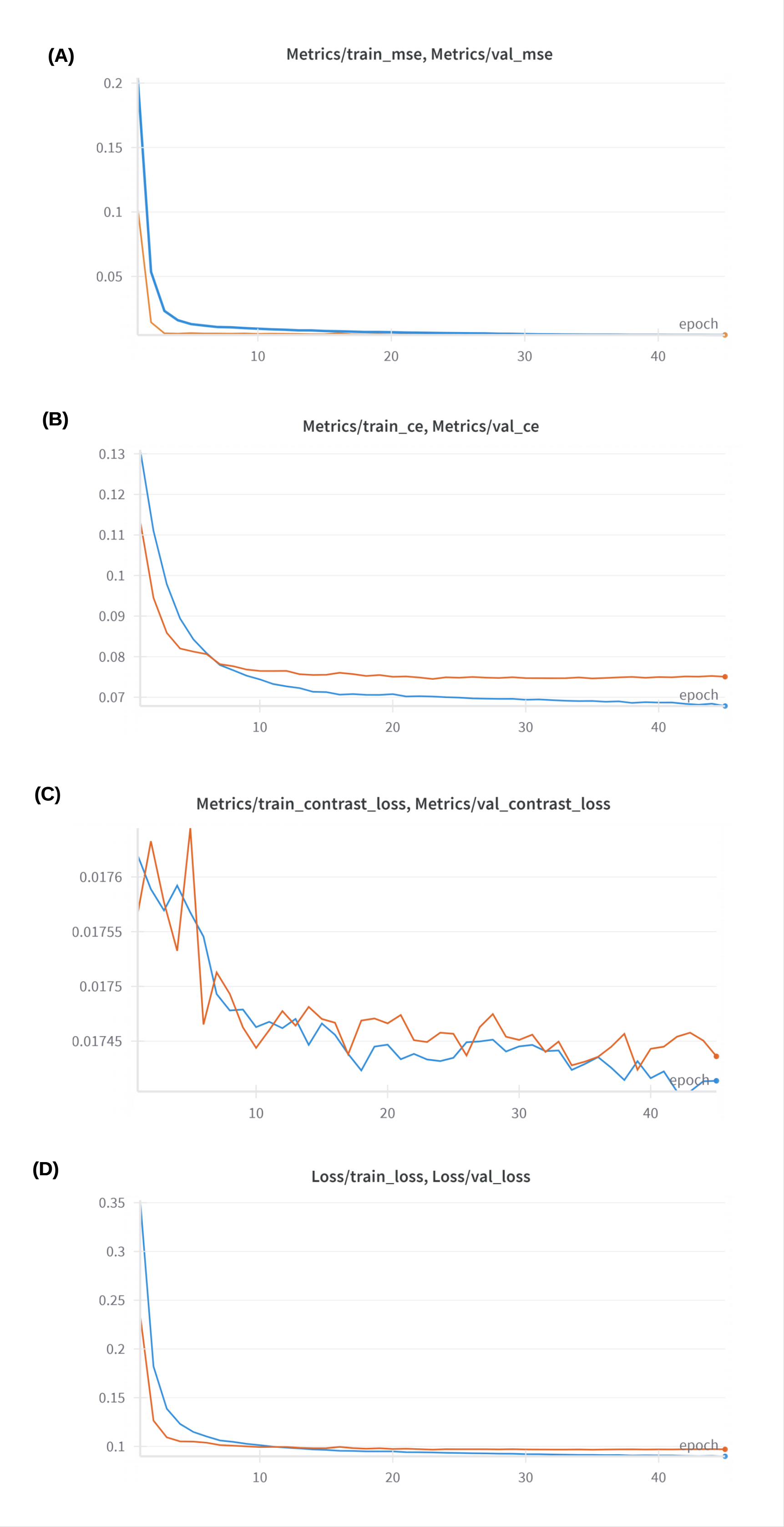


**Fig. S1. (A)**-**(D)** denote MSE Loss, CE Loss, Contrast Loss, and Total Loss, respectively, where the blue one represents the training loss curve and the orange one represents the validation loss curve.

**Table S1. Hyperparameter values chose by ContrastQA. The final parameter values are denoted in bold.**

| Hyperparameter | Search Space |
| --- | --- |
| Weight of $L_{mse}$($w_{mse}$) | **1**,0.9 |
| Weight of $L_{ce}$($w_{ce}$) | **0.1**,0.05,0.01 |
| Weight of $L_{cl}$($w_{cl}$) | 0.1,0.05,0.01,**0.005**,0.001 |
| Learning Rate | 0.005,**0.001**,0.0005,0.0002,0.0001 |
| Weight Decay Rate | 0.002,0.001,**0.0005**,0.0001 |
| Read-Out Module Dropout Rate | 0.1,0.2,0.3,**0.5** |
| Graph Pooling Operator | **Mean**, Max, Sum |
| GVP Hidden Dimension | 128,256,**512** |
| GVP-GNN Layers Number | 1,**2**,3 |
| Temperature Coefficient | **0.07**,0.1,0.2,0.5 |

**Table S2. TMscore ranking loss on each target of the CASP16 dataset.**

| target | ContrastQA | GuijunLab-QA | GuijunLab-PAthreader | MULTICOM | MULTICOM_GATE | MULTICOM_LLM | ModFOLDdock2 | MQA_base |
|---|---|---|---|---|---|---|---|---|
| H1202 | 0.017 | 0.013 | **0.006** | - | 0.024 | 0.013 | 0.008 | 0.012 |
| H1204 | 0.318 | 0.321 | 0.349 | - | 0.284 | **0** | 0.338 | 0.305 |
| H1208 | 0.006 | 0.006 | 0.011 | 0.007 | **0.003** | 0.008 | 0.007 | 0.013 |
| H1213 | 0.021 | 0.045 | 0.017 | **0.004** | 0.021 | 0.019 | 0.043 | 0.017 |
| H1215 | 0.100 | 0.375 | 0.375 | 0.007 | 0.003 | 0.377 | **0.002** | 0.377 |
| H1217 | **0.020** | 0.140 | 0.140 | 0.089 | 0.089 | 0.104 | 0.021 | 0.116 |
| H1220 | 0.038 | **0.034** | 0.038 | 0.038 | 0.043 | 0.035 | **0.034** | 0.127 |
| H1222 | 0.016 | 0.011 | 0.010 | 0.018 | **0.009** | **0.009** | 0.011 | 0.035 |
| H1223 | **0.041** | 0.239 | 0.243 | 0.197 | 0.217 | 0.215 | 0.225 | 0.042 |
| H1225 | 0.019 | 0.026 | 0.022 | **0.012** | 0.019 | 0.016 | 0.022 | 0.018 |
| H1227 | **0.001** | 0.070 | 0.070 | 0.147 | 0.158 | 0.154 | 0.157 | 0.075 |
| H1232 | 0.364 | 0.372 | 0.389 | 0.365 | 0.393 | **0.358** | 0.368 | 0.364 |
| H1233 | **0.002** | 0.003 | 0.004 | 0.008 | 0.008 | 0.003 | **0.002** | 0.004 |
| H1236 | 0.124 | 0.120 | 0.137 | 0.118 | 0.115 | **0.114** | 0.122 | 0.142 |
| H1244 | **0.030** | 0.049 | 0.051 | 0.045 | 0.075 | 0.054 | 0.132 | 0.078 |
| H1245 | 0.310 | 0.113 | 0.117 | 0.129 | 0.078 | 0.076 | **0.011** | 0.124 |
| H1258 | 0.082 | **0.026** | **0.026** | 0.029 | 0.063 | 0.058 | 0.031 | 0.05 |
| H1265 | 0.462 | 0.466 | 0.466 | 0.482 | 0.482 | **0.273** | 0.479 | 0.385 |
| H1267 | 0.535 | **0.508** | 0.510 | 0.509 | 0.515 | **0.508** | **0.508** | 0.511 |
| H1272 | **0.053** | 0.120 | 0.120 | 0.534 | 0.105 | 0.110 | 0.119 | 0.174 |
| T1201o | 0.011 | 0.013 | 0.027 | - | 0.014 | 0.014 | 0.01 | **0.007** |
| T1206o | 0.006 | 0.002 | **0** | 0.001 | 0.014 | 0.001 | 0.041 | 0.006 |
| T1218o | 0.475 | 0.584 | **0.117** | 0.656 | 0.618 | 0.635 | 0.634 | 0.127 |
| T1219o | **0.052** | 0.152 | 0.089 | 0.053 | 0.053 | 0.093 | 0.089 | 0.108 |
| T1234o | 0.040 | 0.041 | 0.614 | 0.072 | 0.048 | 0.038 | 0.606 | **0.034** |
| T1235o | 0.025 | 0.088 | 0.080 | 0.083 | 0.088 | 0.084 | 0.024 | **0.001** |
| T1237o | 0.009 | 0.011 | 0.009 | 0.015 | 0.008 | **0.007** | 0.008 | 0.008 |
| T1240o | 0.380 | 0.326 | 0.655 | 0.383 | 0.386 | 0.357 | **0.236** | 0.386 |
| T1249o | **0.021** | 0.151 | 0.027 | 0.031 | 0.031 | 0.082 | 0.117 | 0.025 |
| T1257o | 0.018 | 0.005 | 0.005 | **0.002** | 0.005 | 0.017 | 0.015 | 0.731 |
| T1259o | 0.543 | 0.012 | 0.028 | **0.007** | 0.538 | 0.543 | 0.542 | 0.544 |
| T1269o | 0.047 | 0.027 | 0.027 | 0.020 | 0.020 | **0** | 0.041 | 0.047 |
| T1270o | 0.213 | 0.210 | 0.227 | 0.202 | 0.196 | 0.206 | **0.136** | 0.218 |
| T1292o | 0.025 | 0.004 | 0.003 | 0.006 | 0.004 | 0.004 | **0.001** | 0.024 |
| T1294o | 0.006 | 0.011 | 0.010 | 0.011 | **0.003** | 0.004 | 0.005 | - |
| T1295o | **0.119** | **0.119** | **0.119** | 0.122 | 0.122 | 0.259 | 0.263 | - |
| T1298o | 0.019 | 0.297 | 0.327 | 0.322 | 0.302 | 0.295 | 0.310 | **0.009** |
| average | **0.123** | 0.138 | 0.148 | 0.139 | 0.139 | 0.139 | 0.155 | 0.150 |

Note: The specific data of each target were intercepted from the CASP official website. Specific H1265, T1219o, and T1295o are based on the results of each team using US-align to get the TMscore. Data in bold indicates the best result.

**Table S3. GDT-TS ranking loss on each target of the CASP16 dataset.**

| target | ContrastQA | GuijunLab-QA | GuijunLab-PAthreader | MULTICOM | MULTICOM_GATE | MULTICOM_LLM | ModFOLDdock2 | MQA_base |
|---|---|---|---|---|---|---|---|---|
| H1202 | 0.062 | 0.044 | **0.022** | - | 0.078 | 0.046 | 0.031 | 0.043 |
| H1204 | 0.281 | 0.282 | 0.278 | - | 0.219 | **0** | 0.275 | 0.283 |
| H1208 | 0.039 | 0.047 | **0.038** | 0.054 | 0.049 | 0.056 | 0.062 | 0.100 |
| H1213 | 0.114 | 0.217 | 0.130 | **0.037** | 0.137 | 0.101 | 0.214 | 0.130 |
| H1215 | 0.179 | 0.365 | 0.365 | 0.026 | 0.011 | 0.369 | **0.006** | 0.368 |
| H1217 | 0.052 | 0.222 | 0.222 | **0.045** | **0.045** | 0.166 | 0.062 | 0.072 |
| H1220 | 0.073 | **0.051** | 0.073 | 0.089 | 0.079 | 0.062 | 0.106 | 0.111 |
| H1222 | **0.032** | 0.059 | 0.044 | 0.079 | 0.052 | 0.052 | 0.059 | 0.134 |
| H1223 | **0.055** | 0.394 | 0.331 | 0.273 | 0.375 | 0.374 | 0.333 | 0.088 |
| H1225 | 0.039 | 0.056 | 0.045 | 0.038 | **0.024** | 0.033 | 0.051 | 0.068 |
| H1227 | **0.062** | 0.202 | 0.202 | 0.129 | 0.119 | 0.143 | 0.195 | 0.205 |
| H1232 | 0.231 | 0.230 | 0.218 | 0.225 | 0.403 | **0.215** | 0.233 | 0.235 |
| H1233 | 0.024 | 0.024 | 0.032 | 0.072 | 0.072 | 0.024 | **0.012** | 0.028 |
| H1236 | 0.089 | 0.097 | 0.075 | **0.038** | 0.069 | 0.063 | 0.098 | 0.159 |
| H1244 | 0.054 | 0.042 | **0.031** | 0.105 | 0.091 | 0.043 | 0.184 | 0.098 |
| H1245 | 0.375 | 0.202 | 0.214 | 0.226 | 0.162 | 0.158 | 0.023 | 0.216 |
| H1258 | 0.020 | 0.017 | 0.017 | 0.069 | 0.027 | **0.007** | 0.070 | 0.045 |
| H1265 | 0.127 | 0.127 | 0.127 | 0.128 | 0.128 | 0.095 | 0.127 | **0.060** |
| H1267 | 0.527 | **0.393** | 0.411 | 0.396 | 0.445 | **0.393** | **0.393** | 0.521 |
| H1272 | **0** | 0.098 | 0.098 | 0.334 | 0.082 | 0.083 | 0.042 | 0.108 |
| T1201o | 0.046 | 0.050 | 0.082 | - | 0.055 | 0.055 | 0.060 | **0.031** |
| T1206o | 0.036 | 0.012 | 0.004 | 0.009 | 0.059 | **0.003** | 0.082 | 0.020 |
| T1218o | 0.363 | 0.400 | 0.239 | 0.423 | 0.421 | 0.429 | 0.412 | **0.222** |
| T1219o | **0.033** | 0.167 | 0.077 | **0.033** | **0.033** | 0.076 | 0.077 | 0.110 |
| T1234o | 0.062 | 0.073 | 0.582 | 0.215 | **0.048** | 0.098 | 0.57 | 0.090 |
| T1235o | 0.135 | 0.183 | 0.135 | 0.149 | 0.182 | 0.150 | 0.156 | **0.013** |
| T1237o | 0.062 | 0.080 | 0.054 | 0.102 | **0.029** | 0.058 | 0.068 | 0.064 |
| T1240o | 0.476 | 0.457 | 0.471 | 0.490 | 0.491 | 0.466 | **0.419** | 0.495 |
| T1249o | **0.078** | 0.383 | 0.123 | 0.165 | 0.161 | 0.303 | 0.379 | 0.108 |
| T1257o | 0.138 | 0.051 | 0.051 | **0.011** | 0.051 | 0.138 | 0.129 | 0.587 |
| T1259o | 0.045 | 0.044 | 0.069 | 0.015 | 0.097 | **0.003** | 0.028 | 0.046 |
| T1269o | 0.080 | 0.033 | 0.033 | 0.102 | 0.102 | **0** | 0.051 | 0.089 |
| T1270o | 0.056 | 0.038 | 0.041 | **0.019** | 0.050 | 0.051 | 0.043 | 0.045 |
| T1292o | 0.028 | 0.005 | 0.005 | 0.009 | 0.006 | 0.006 | **0** | 0.026 |
| T1294o | **0.013** | 0.033 | 0.029 | 0.034 | 0.017 | 0.017 | 0.014 | - |
| T1295o | **0.166** | 0.202 | 0.202 | 0.203 | 0.203 | 0.201 | 0.207 | - |
| T1298o | 0.044 | 0.206 | 0.091 | 0.266 | 0.218 | 0.210 | 0.252 | **0.037** |
| average | **0.116** | 0.151 | 0.142 | 0.136 | 0.132 | 0.128 | 0.149 | 0.144 |

Note: The specific data of each target were intercepted from the CASP official website. Specific H1265, T1219o, and T1295o are based on the results of each team using openstructure to get the GDT-TS. Data in bold indicates the best result.

**Table S4. TMscore ranking loss of easy targets on the CASP16 dataset.**

| target | ContrastQA | GuijunLab-QA* | GuijunLab-PAthreader* | MULTICOM_LLM* | ModFOLDdock2* | MQA_base* | TopoQA |
|---|---|---|---|---|---|---|---|
| T1206o | 0.006 | 0.002 | 0 | 0.001 | 0.041 | 0.006 | 0.005 |
| T1234o | 0.040 | 0.041 | 0.614 | 0.038 | 0.606 | 0.034 | 0.044 |
| T1235o | 0.025 | 0.088 | 0.080 | 0.084 | 0.024 | 0.001 | 0.087 |
| T1259o | 0.543 | 0.012 | 0.028 | 0.543 | 0.542 | 0.544 | 0 |
| T1292o | 0.025 | 0.004 | 0.003 | 0.004 | 0.001 | 0.024 | 0.004 |
| T1294o | 0.006 | 0.011 | 0.010 | 0.004 | 0.005 | - | 0.009 |
| average | 0.108 | 0.026 | 0.123 | 0.112 | 0.203 | 0.122 | **0.025** |

Note: Data in bold indicates the best result. Those with * refer to the methods of the teams participating in CASP16, where the data are extracted from the official website results.

**Table S5. Top scores mean on the CASP16 dataset.**

| Method | TMscore | | GDT-TS | |
|---|---|---|---|---|
| | Top3 mean (↑) | Top5 mean (↑) | Top3 mean (↑) | Top5 mean (↑) |
| GNN-DOVE | 0.599 | 0.605 | 0.469 | 0.470 |
| DProQA | 0.709 | 0.706 | 0.530 | 0.546 |
| ComplexQA | 0.574 | 0.582 | 0.406 | 0.414 |
| VoroIF-GNN | 0.709 | 0.715 | 0.556 | 0.572 |
| TopoQA | 0.698 | 0.703 | 0.558 | 0.568 |
| ContrastQA | **0.737** | **0.725** | **0.609** | **0.593** |

Note: Data in bold indicates the best result. Top3 mean and Top5 mean are the average of the true scores of the top three and five predicted decoy structures, respectively, the larger the value indicates the higher performance the method has.

**Table S6. TMscore ranking loss on each target of the ABAG-AF3 dataset.**

| Target | ContrastQA | ComplexQA | DProQA | GNN-DOVE | TopoQA | VoroIF-GNN |
|---|---|---|---|---|---|---|
| 7o9w | 0.045 | 0.069 | 0.034 | 0.069 | **0.018** | 0.039 |
| 7om4 | 0.060 | 0.169 | 0.168 | 0.168 | 0.169 | **0.033** |
| 7r40 | 0.018 | 0.021 | **0.015** | 0.021 | 0.022 | 0.017 |
| 7s0e | **0.018** | 0.021 | 0.035 | 0.073 | 0.024 | **0.018** |
| 7sbg | 0.020 | 0.197 | 0.134 | 0.134 | **0.007** | 0.022 |
| 7sd3 | 0.085 | 0.087 | 0.082 | **0.023** | 0.090 | 0.031 |
| 7sgm | 0.019 | 0.016 | 0.016 | 0.020 | 0.016 | **0** |
| 7sjn | 0.015 | 0.010 | 0.006 | **0** | 0.015 | 0.007 |
| 7sjo | 0.056 | 0.058 | **0.008** | **0.008** | 0.053 | 0.053 |
| 7su0 | **0** | 0.059 | 0.030 | **0** | 0.036 | **0** |
| 7su1 | **0** | 0.003 | 0.001 | 0.007 | 0.008 | 0.008 |
| 7swn | 0.033 | 0.183 | 0.002 | 0.372 | **0** | 0.014 |
| 7t25 | 0.015 | 0.020 | 0.036 | 0.012 | 0.036 | **0.012** |
| 7t73 | 0.012 | 0.008 | 0.007 | 0.007 | 0.011 | **0.006** |
| 7t77 | 0.008 | 0.009 | 0.008 | 0.007 | 0.009 | **0.004** |
| 7tee | **0.020** | 0.113 | 0.110 | 0.102 | 0.036 | 0.024 |
| 7tfo | 0.019 | 0.012 | 0.007 | 0.013 | **0.003** | 0.019 |
| 7tyv | 0.037 | 0.028 | **0** | 0.028 | 0.035 | 0.046 |
| 7ued | 0.049 | **0** | 0.050 | 0.051 | 0.050 | 0.051 |
| 7vng | 0.004 | 0.176 | 0.186 | **0** | 0.009 | 0.005 |
| 7vyr | 0.007 | 0.004 | 0.004 | 0.117 | 0.005 | **0.001** |
| 7wo5 | 0.013 | 0.020 | **0.009** | **0.009** | 0.018 | 0.020 |
| 7wrv | **0.062** | 0.293 | 0.297 | 0.319 | 0.319 | 0.319 |
| 7x7o | **0** | **0** | **0** | **0** | **0** | **0** |
| 7yqx | 0.002 | **0.001** | 0.002 | **0.001** | 0.004 | 0.002 |
| 7yqz | 0.018 | **0.015** | 0.020 | 0.016 | 0.018 | 0.016 |
| 7z2m | 0.014 | **0** | 0.013 | **0** | 0.013 | 0.002 |
| 7z4t | 0.004 | 0.006 | 0.153 | 0.196 | 0.007 | **0.003** |
| 7zf9 | 0.017 | 0.001 | 0.008 | **0** | 0.009 | 0.031 |
| 7zr7 | 0.016 | 0.016 | **0** | 0.002 | 0.001 | 0.017 |
| 8b7w | 0.145 | 0.169 | 0.169 | 0.113 | **0.028** | 0.145 |
| 8f8x | 0.032 | 0.033 | 0.033 | 0.032 | 0.033 | **0** |
| 8gv6 | 0.042 | 0.036 | 0.041 | 0.041 | 0.043 | **0.011** |
| 8gv7 | **0.052** | **0.052** | 0.132 | 0.114 | 0.090 | 0.113 |
| 8gz5 | **0.004** | 0.026 | 0.337 | 0.007 | 0.059 | 0.009 |
| **average** | **0.028** | 0.055 | 0.062 | 0.060 | 0.037 | 0.031 |

Note: Data in bold indicates the best result. In the ranking loss section, a target may have multiple top1-ranked models, and we took the average of the ranking losses of these models as the ranking loss for this target.

**Table S7. GDT-TS ranking loss on each target of the ABAG-AF3 dataset.**

| Target | ContrastQA | ComplexQA | DProQA | GNN-DOVE | TopoQA | VoroIF-GNN |
|---|---|---|---|---|---|---|
| 7o9w | 0.140 | 0.161 | 0.095 | 0.161 | **0.089** | 0.104 |
| 7om4 | 0.043 | 0.097 | 0.103 | 0.094 | 0.097 | **0.028** |
| 7r40 | **0** | 0.001 | 0.001 | 0.001 | 0.001 | 0.001 |
| 7s0e | 0.015 | 0.025 | **0.005** | 0.009 | **0.005** | 0.015 |
| 7sbg | 0.016 | 0.129 | **0.008** | **0.008** | 0.013 | 0.010 |
| 7sd3 | 0.034 | 0.030 | 0.033 | 0.031 | **0.029** | 0.034 |
| 7sgm | 0.024 | 0.009 | 0.017 | 0.006 | **0** | 0.016 |
| 7sjn | 0.009 | **0.005** | 0.007 | 0.010 | 0.009 | **0.005** |
| 7sjo | **0.001** | **0.001** | **0.001** | **0.001** | **0.001** | **0.001** |
| 7su0 | 0.001 | 0.084 | 0.104 | **0** | 0.098 | 0.001 |
| 7su1 | **0** | 0.017 | 0.007 | 0.046 | 0.055 | 0.055 |
| 7swn | 0.079 | 0.187 | 0.001 | 0.312 | **0** | 0.034 |
| 7t25 | **0.008** | 0.013 | 0.016 | 0.023 | 0.030 | 0.019 |
| 7t73 | 0.009 | **0.003** | 0.021 | **0.003** | **0.003** | 0.022 |
| 7t77 | 0.008 | 0.047 | 0.011 | **0.005** | 0.047 | **0.005** |
| 7tee | 0.005 | 0.007 | 0.004 | 0.003 | 0.006 | **0.002** |
| 7tfo | 0.006 | 0.009 | 0.018 | 0.005 | **0.003** | 0.006 |
| 7tyv | 0.010 | 0.007 | 0.010 | **0.003** | 0.011 | 0.012 |
| 7ued | **0.005** | 0.025 | 0.007 | 0.010 | 0.006 | 0.011 |
| 7vng | 0.008 | 0.183 | 0.068 | 0.004 | 0.038 | **0** |
| 7vyr | 0.002 | **0** | **0** | 0.002 | 0.002 | 0.002 |
| 7wo5 | 0.005 | **0.003** | 0.005 | 0.005 | 0.006 | 0.005 |
| 7wrv | **0.003** | 0.053 | 0.055 | 0.216 | 0.216 | 0.215 |
| 7x7o | 0.001 | 0.001 | 0.001 | 0.001 | **0** | 0.001 |
| 7yqx | **0** | 0.003 | 0.003 | 0.003 | 0.003 | 0.003 |
| 7yqz | 0.001 | 0.002 | 0.001 | **0** | 0.001 | **0** |
| 7z2m | 0.024 | **0** | 0.024 | **0** | 0.024 | 0.003 |
| 7z4t | **0.009** | **0.009** | 0.238 | 0.259 | 0.025 | 0.026 |
| 7zf9 | 0.063 | 0.010 | 0.035 | **0.002** | 0.041 | 0.102 |
| 7zr7 | 0.006 | 0.006 | 0.006 | 0.005 | 0.004 | **0.003** |
| 8b7w | 0.145 | 0.137 | 0.137 | 0.131 | **0** | 0.155 |
| 8f8x | 0.003 | 0.003 | 0.003 | **0.001** | 0.002 | 0.003 |
| 8gv6 | **0.005** | 0.013 | 0.007 | 0.007 | 0.008 | 0.011 |
| 8gv7 | 0.062 | 0.062 | 0.091 | 0.018 | **0.013** | 0.024 |
| 8gz5 | 0.002 | **0** | 0.013 | 0.002 | 0.012 | 0.003 |
| **average** | **0.021** | 0.038 | 0.033 | 0.040 | 0.026 | 0.027 |

Note: Data in bold indicates the best result. In the ranking loss section, a target may have multiple top1-ranked models, and we took the average of the ranking losses of these models as the ranking loss for this target.

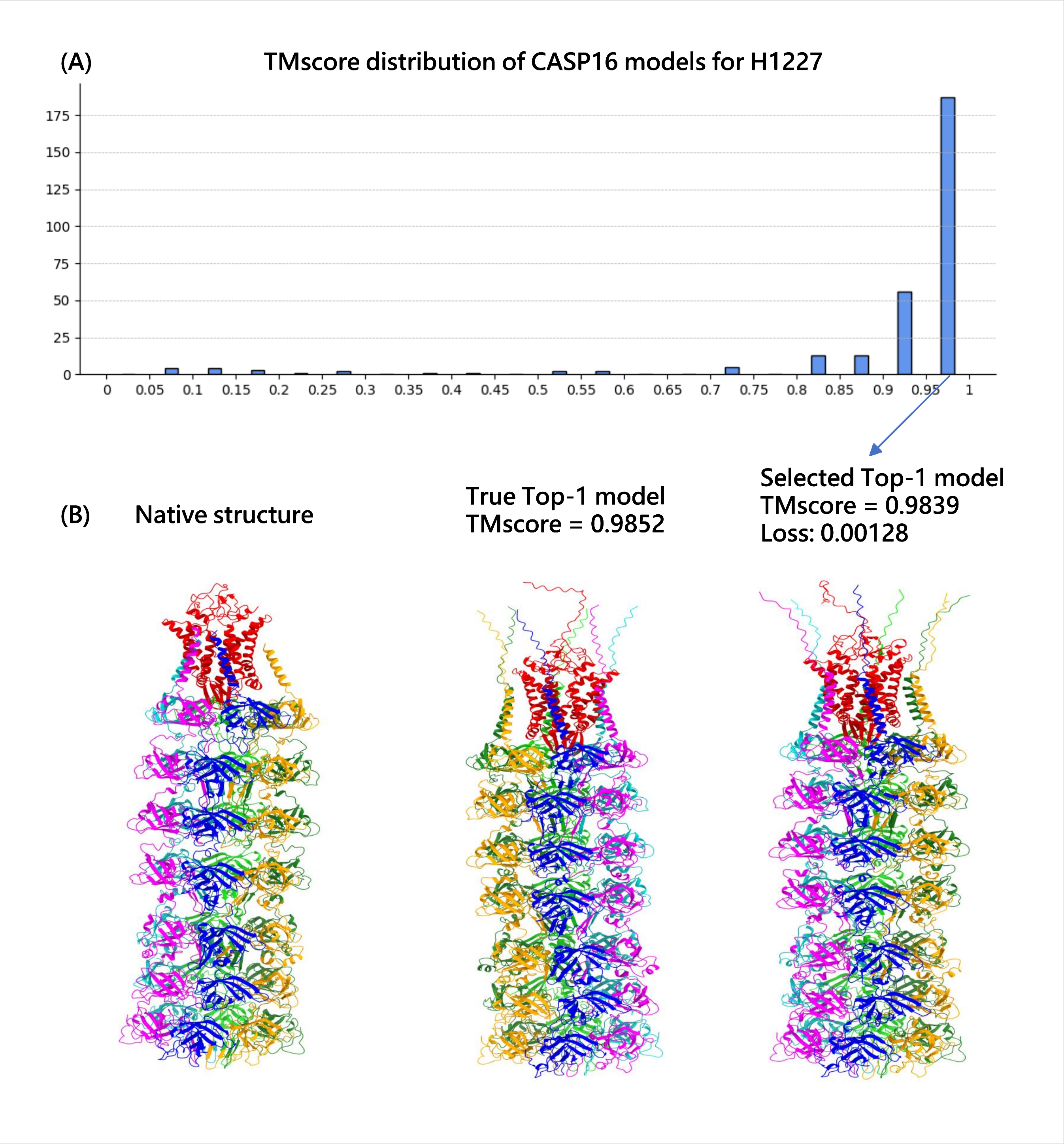


**Fig. S2.** **(A)** is the TMscore distribution of CASP16 models for H1227. **(B)** includes corresponding native structure, the true TOP-1 model (TMscore is 0.985) and the ContrastQA Top-1 selected model (TMscore is 0.984, ContrastQA achieved a 0.00128 TMscore ranking loss.) to target H1227.
